# Can a population targeted by a CRISPR-based homing gene drive be rescued?

**DOI:** 10.1101/2020.03.17.995829

**Authors:** Nicolas O. Rode, Virginie Courtier-Orgogozo, Florence Débarre

## Abstract

CRISPR-based homing gene drive is a genetic control technique aiming to modify or eradicate natural populations. This technique is based on the release of individuals carrying an engineered piece of DNA that can be preferentially inherited by the progeny. Developing countermeasures is important to control the spread of gene drives, should they result in unanticipated damages. One proposed countermeasure is the introduction of individuals carrying a brake construct that targets and inactivates the drive allele but leaves the wild-type allele unaffected. Here we develop models to investigate the efficiency of such brakes. We consider a variable population size and use a combination of analytical and numerical methods to determine the conditions where a brake can prevent the extinction of a population targeted by an eradication drive. We find that a brake is not guaranteed to prevent eradication and that characteristics of both the brake and the drive affect the likelihood of recovering the wild-type population. In particular, brakes that restore fitness are more efficient than brakes that do not. Our model also indicates that threshold-dependent drives (drives that can spread only when introduced above a threshold) are more amenable to control with a brake than drives that can spread from an arbitrary low introduction frequency (threshold-independent drives). Based on our results, we provide practical recommendations and discuss safety issues.

**Article summary for Issue Highlights:** Homing gene drive is a new genetic control technology that aims to spread a genetically engineered DNA construct within natural populations even when it impairs fitness. In case of unanticipated damages, it has been proposed to stop homing gene drives by releasing individuals carrying a genedrive brake; however, the efficiency of such brakes has been little studied. The authors develop a model to investigate the dynamics of a population targeted by a homing drive in absence or in presence of brake. The model provides insights for the design of more efficient brakes and safer gene drives.

## Introduction

The use of engineered gene drives has been proposed as a technique for population control with potential applications in public health, agriculture and conservation (Burt 2003; Esvelt *et al*. 2014). This technique relies on the release of genetically engineered individuals that can rapidly propagate a transgene of interest into wild populations. Gene drive can be designed to modify, suppress or eradicate various target species (Scott *et al*. 2018; Rode *et al*. 2019). Potential target species include disease vectors (e.g. *Anopheles gambiae*, the main vector of malaria in Africa; Kyrou *et al*. 2018), agricultural pests (e.g. *Drosophila suzukii*, a major pest of soft fruits; Scott *et al*. 2018) or invasive rodents (e.g. invasive house mouse or black rats that threaten biodiversity on islands; Leitschuh *et al*. 2018).

Due to the universality of CRISPR genome editing, CRISPR-based gene drives can potentially be applied to a wide variety of organisms (Esvelt *et al*. 2014; Raban *et al*. 2020). Diverse CRISPR-based gene drive systems have already been developed in the laboratory as proofs-of-principle in a few model organisms (homing, split homing, translocation, X-shredder, killer-rescue, cleave-and-rescue and TARE gene drives; Webster *et al*. 2019; see Raban *et al*. 2020 for a review; Champer *et al*. 2020) or as theoretical possibilities (daisy chain drives; Noble *et al*. 2019). Gene drives have so far only been tested in the laboratory and no field trial has been conducted yet.

Among these systems, CRISPR-based homing gene drives are the most adaptable to new species and populations and the most advanced in terms of technological development (Raban *et al*. 2020). They involve a piece of DNA that includes a guide RNA (gRNA) gene and a *cas9* gene (encoding the Cas9 endonuclease). The gRNA is designed to recognize a specific sequence in a wild-type chromosome, so that that in heterozygotes carrying a drive allele and a wild-type allele, the Cas9-gRNA molecular complex will cut the wild-type chromosome at the target site. The resulting double-strand DNA break can then be repaired through homology-directed repair (also known as “gene conversion”), using the drive allele as a template, which is designed to harbor sequences identical to the ones flanking the target site. Consequently, the drive allele is transmitted to the next generation at rates beyond those of regular Mendelian inheritance and, if its parameters allow it, will rapidly spread within the target population.

Homing gene drives are sometimes considered as “threshold-independent drives”, i.e. as being able to spread in a population from an arbitrary low introduction frequency (e.g. Marshall and Akbari 2018). Mathematical models of homing gene drives (e.g. Deredec *et al*. 2008; Alphey and Bonsall 2014; Unckless *et al*. 2015; Tanaka *et al*. 2017) have however shown that depending on various parameters (the efficacy of gene conversion, its timing, the fitness cost incurred by the drive allele and its dominance over the wild-type allele), some of the homing gene drives can be threshold-dependent, i.e. only spread if they are introduced above a threshold frequency. Mathematically, when there is an equilibrium at an intermediate frequency of the drive allele (0 < *p_D_* < 1) and when this equilibrium is unstable, then the drive is threshold-dependent; the value of the drive allele frequency at this equilibrium is the threshold above which the drive has to be introduced to spread (Deredec *et al*. 2008).

Given that gene drives can potentially impact biodiversity, national sovereignty and food security (Oye *et al*. 2014; Akbari *et al*. 2015; DiCarlo *et al*. 2015; NASEM 2016; Montenegro de Wit 2019), there is a crucial need to develop strategies to minimize the risks of unintentional spread (e.g. following the escape of gene drive individuals from a laboratory) and to mitigate unanticipated or premeditated and malevolent harm to humans or the environment. For example, a CRISPR-based eradication drive may spread into a non-target population or species (Noble *et al*. 2018; Courtier-Orgogozo *et al*. 2019a; Rode *et al*. 2019); a modification drive may alter the target population in an unexpected, detrimental manner; or a gene drive could be used as bioweapon (Gurwitz 2014). Decreasing the environmental risks associated with the development of this technology can be achieved by designing safer gene drives whose spread can be controlled spatially or temporally (Marshall and Akbari 2018; Raban *et al*. 2020) and by developing countermeasures to stop the spread of an ongoing gene drive (Esvelt *et al*. 2014; Gantz and Bier 2016; Vella *et al*. 2017).

Several countermeasure strategies for CRISPR-based homing gene drives have been proposed. One strategy is to use gene drives whose non-Mendelian transmission is conditional on the presence of synthetic molecules in the environment of the target species, so that the removal of the synthetic molecule is expected to stop the spread of the gene drive, and natural selection to remove the drive from the population (Esvelt *et al*. 2014; Del Amo *et al*. 2020). However, the development of such molecule-dependent drives is still at its infancy and may have to be tailored for each ecosystem and target species. Another strategy is to introduce resistant individuals carrying a modified target locus that prevents homing (“synthetic resistant” (SR) allele; Burt 2003; Champer *et al*. 2016; Vella *et al*. 2017). However, this strategy results in a modified population with 100% resistant individuals and does not allow the recovery of the original wild-type population. In addition, synthetic resistant alleles are predicted to be rather ineffective against replacement drives with small fitness costs (Vella *et al*. 2017), because of the limited selective advantage of synthetic resistant alleles. Alternatively, it has been proposed to release suppressor individuals that carry a new piece of DNA which will eventually lead to the knock-out of the initial gene drive (Esvelt *et al*. 2014; Marshall and Akbari 2018). These alternative countermeasures rely on gene conversion and can be used against virtually any type of CRISPR-based homing gene drive. Two types can be distinguished. The first type are countermeasures that include the *cas9* gene and that can target either the drive allele only (reversal drives sensu Esvelt *et al*. 2014; overwriting drives; DiCarlo *et al*. 2015) or both the drive and wild-type alleles (immunizing reversal drive (IRD); Esvelt *et al*. 2014; Vella *et al*. 2017). However, with these strategies, a functional *cas9* gene will remain in the final population, which may increase the risk of subsequent genetic modifications such as translocations, and possible negative environmental outcomes (Courtier-Orgogozo *et al*. 2019b). The second type are countermeasures that do not encode *cas9* and rely instead on the *cas9* gene present in the initial gene drive construct. They can be contained in a single locus (ERACR: element for reversing the autocatalytic chain reaction, Gantz and Bier 2016; CATCHA: Cas9-triggered chain ablation, Wu *et al*. 2016), or be across two loci (CHACR: construct hitchhiking on the autocatalytic chain reaction, Gantz and Bier 2016). These countermeasures might be safer for the environment, due to the absence of a functional *cas9* gene. To our knowledge, only the CATCHA brakes have been implemented in the lab (Supplemental Material, Figure S1); CHACR may be slow to spread due to its two-locus structure, while ERACR may be sensitive to the evolution of resistance at its target sites (*cas9*-flanking sequences whose mutation does not affect enzyme function).

We focus here on the -- in our opinion -- best gene-drive-based countermeasures proposed so far, the cas9-devoid reversal drives (CATCHA, ERACR), which we call hereafter “brakes” for simplicity. In drive/brake heterozygotes, the encoded guide RNA(s) target and inactivate the *cas9* gene of the initial gene drive construct. Such brakes should be especially efficient, because even in absence of homology-directed repair, the drive’s *cas9* gene (targeted by the brake) is expected to be inactivated. However, for simplicity, we will not model this additional scenario here.

Although mathematical modelling of the effects of brakes has been recommended (Wu *et al*. 2016), to our knowledge only two such studies have been published (Vella *et al*. 2017; Girardin *et al*. 2019). Vella et al. found that the introduction of a brake leads to a polymorphic equilibrium with transient oscillatory dynamics (Figure 2d,e in Vella *et al*. 2017). They also showed that brakes with smaller fitness costs increased the likelihood of long-term eradication of the homing gene drive (Figure 3 in Vella *et al*. 2017). We note that because Vella *et al*. (2017) assumed 100% cleavage and germline conversion, the drive they modeled was threshold-independent (Deredec *et al*. 2008). Girardin et *al*. (2019) considered a spatial model, and found that a brake could stop a spatially spreading drive only if the drive was threshold-dependent, and that threshold-independent drives led to an infinite spatial chase of the drive by the brake. While both studies provided insights on our ability to control an ongoing gene drive, they had limitations. First, Vella *et al*. (2017) used classical population-genetic frameworks, and focused on allele frequency dynamics, ignoring changes in population size. Changes in total population size were also not the focus of Girardin et *al*. (2019). Both studies omitted potential demographic feedbacks on allele frequency changes, which are likely to be important for eradication drives. It thus remains unknown whether a brake can prevent the extinction of a population targeted by an eradication drive. Second, both studies used deterministic models. Vella et al. acknowledged that oscillations of the allele frequencies in their model could lead to the stochastic loss of an allele. Similar oscillations were observed by Girardin et *al*. (2019), but their implications were not explored.

To address some of the limitations of previous models and examine further the effectiveness of brakes, we model here the dynamics of a population targeted by a drive, into which brake-carrying individuals are released. We consider a variable population size and its potential feedback onto gene frequency changes, and we also develop a stochastic version of the model. We compare two timings of gene conversion for gene drive and brake alleles (in the germline or zygote, Figure 1) and explore the role of parameters such as level of dominance, cleavage efficiency, brake-associated fitness costs (whether or not it restores fitness), and the type of fitness component targeted by the gene drive (embryo survival, fecundity or adult death rate). We contrast brakes that restore fitness with those that do not. Implementing brakes that restore fitness (i.e. “specific brakes”) require prior knowledge of the gene disrupted by the homing drive in order to include in the brake a recoded version of this gene along with a gRNA that targets the *cas9* sequence of the drive allele. With brakes that restore fitness, drive-brake heterozygous individuals have higher fitness than drive homozygotes, but may have lower fitness than wild-type homozygotes (as they may incur a small fitness cost due to the expression of the gRNA). Implementing CATCHA brakes that do not restore fitness (i.e. “universal brakes”) does not require prior knowledge of the gene disrupted by the homing drive, because such brakes only include a gRNA that targets the *cas9* sequence of the drive allele. With brakes that do not restore fitness, drive-brake heterozygous individuals have the same fitness as drive homozygotes.

**Figure 1:**
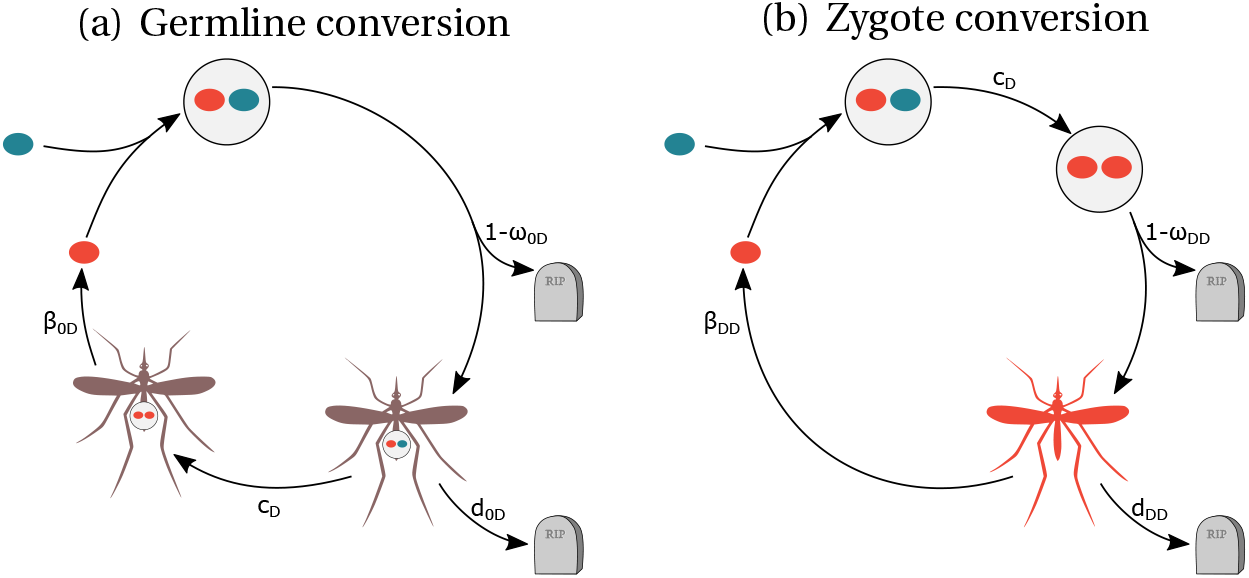
Life-cycles with the two timings of gene conversion, germline (a) and zygote (b). The blue color corresponds to the wild-type allele, the red color to the drive allele and drive-homozygous individuals; the drive/wild- type heterozygous individual is represented in purple. The tombstone represents death. Notation: 0: WT, D: drive; *c* probability of gene conversion; *ω:* zygote survival; *d*: adult mortality; *β*: adult fecundity.

Eradication drives currently under development target genes involved in female development in various human-disease vectors (Kyrou *et al*. 2018) or agricultural pests (Li and Scott 2016). These drives are threshold-independent and pose the greatest risks of unwanted spread. We focus on this type of eradication drives in the numerical part of our study. We aim at finding the characteristics of the brakes that can efficiently stop an ongoing gene drive and allow the recovery of a wild-type population.

## Methods

### Analytical model

With three different alleles in the population (wild-type 0, drive *D* and brake *B*), we need to follow the dynamics of six diploid genotypes. We denote by *G* = {00, 0*D*, *DD*, 0*B*, *DB*, *BB*}the set of all possible genotypes. To take into account gene drives that affect population size (as do e.g. eradication drives), we consider the densities of individuals of each genotype and do not focus solely on genotype frequencies as previous models did (Deredec *et al*. 2008; Unckless *et al*. 2015; Vella *et al*. 2017; Girardin *et al*. 2019). We denote the density of individuals of genotype *g* by *N_g_* and the total population density by A (omitting the time dependence (*t*) for concision; *N* = ∑_*g*_ *N_g_*). We consider three traits affecting fitness that can vary among genotypes: the survival of zygotes (*ω_g_*), the death rate of adults (*d_g_*), and the fecundity of adults (*β_g_*). We assume that reproduction is density-dependent: it depends on the total population size *N*, following a classical logistic regulation with carrying capacity *K*. The death rate, on the other hand, is density-independent. The change over time in the density of individuals of genotype *g* is given by

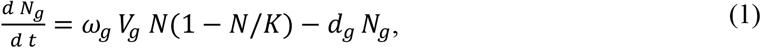

where *V_g_* corresponds to the production of new individuals of genotype *g* through sexual reproduction and depends on the abundances of all genotypes, their fecundities *β_g_*, but also on the timing of gene conversion. The formulas of the *V_g_* terms for each timing of gene conversion are given in the Appendix (and also provided in the supplementary Mathematica file).

We consider that gene conversion in 0*D* or Dβheterozygous individuals can either occur in the germline or in the zygote (Figure 1). When gene conversion occurs in the germline, 0*D* and *DB* heterozygous individuals produce more than 50% of *D* and *B* gametes respectively. When gene conversion occurs in newly formed zygotes (i.e. immediately after fertilization), 0*D* and *DB* heterozygous individuals are converted into *DD* and *BB* homozygotes respectively and have the corresponding traits. For both types of gene conversion, we denote the probabilities of successful gene conversion by drive and by brake alleles by *c_D_* and *c_B_* respectively.

### Numerical explorations

While our analytic results are obtained with generic parameters, numerical explorations require specific parameter values. The number of parameter combinations to explore being very vast, we make a few assumptions to reduce it. First, we consider that drive and brake affect either (i) zygote survival (*ω*), (ii) adult survival (*d*) or (iii) adult fecundity (*β*), all other parameters remaining equal across genotypes. To model an eradication drive, we chose *ω_DD_*, *d_DD_* or *β_DD_* such that a 100% drive population is not viable, and we standardised the parameters to yield the same negative equilibrium value of population size (specifically, we set 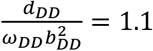, see Table S3 and Mathematica Appendix for details). We consider that either the brake allele does not restore the fitness loss due to the drive allele (i.e. it has the same fitness as the drive allele), or that the brake allele restores partially the fitness loss and has a small fitness cost compared to the wild-type allele (i.e. it contains a specific cargo that helps to restore fitness). We use the same dominance parameter, *h*, for both drive and brake alleles. This choice is justified both when the brake restores and when it does not restore fitness (see the Appendix). For juvenile survival, the parameters of heterozygotes therefore read:

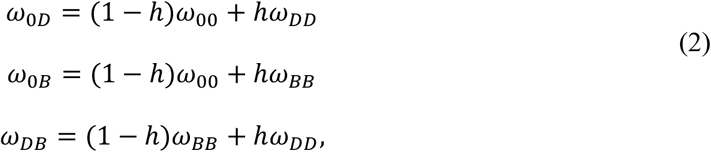

and likewise for *d* and *β* parameters. In the numerical part of the study, we consider either complete recessivity (*h* = 0) or codominance (*h* = 0.5).

We have 24 combinations of parameters (2 timings of gene conversion x 3 traits affected x dominance values x 2 types of brake). For each of them, we consider different timings of introduction of the brake in the population; the timing is given in terms of the current frequency *f_I_* of the drive allele in the population at the time at which the brake is introduced. The *N*^(0)^_0*B*_ parameter represents the number of released wild-type/brake heterozygous individuals. Unless stated, we assume that *N*^(0)^_0*B*_ = 100. Other parameters are shown in tables S1–S3.

### Reformulating the model

Our model is initially defined in terms of genotype densities (equation 1). To simplify the analyses, we reparametrize the model in terms of total population size *N*, allele frequencies *p_D_* and *p_B_* (we have *p*_0_ = 1 – *p_D_* – *p_B_*), and deviations from Hardy-Weinberg for each of the three heterozygotes (*δ*_0*D*_, *δ*_0*B*_, *δ*_*DB*_):

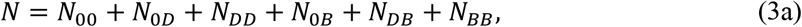

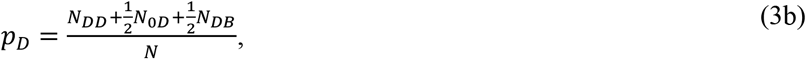

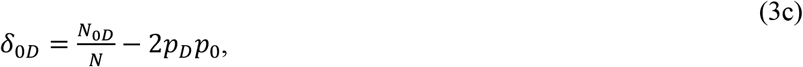

and likewise for *p_B_*, *δ*_0*B*_ and *δ_DB_* (the full equations are calculated in the supplementary Mathematica file).

As usual with most continuous-time models (Nagylaki and Crow 1974), we cannot neglect deviations from Hardy-Weinberg frequencies here (unlike models with discrete, non-overlapping generations). The reformulated model (system (3)) also highlights interactions between total population size *N* and changes in allele frequencies (i.e., eco-evolutionary feedbacks). The population growth rate depends on population composition, since fecundity or survival parameters are genotype-dependent. Reciprocally, changes in allele frequencies depend on the size of the population. This is because gene conversion, which modifies allele frequencies, takes place upon reproduction (either in the germline, or in the newly formed zygote). Given that reproduction is negatively density-dependent, changes in the frequencies of drive and brake alleles slow down when population size is larger.

### Stability analyses

We use the reformulated version of the model (system (3)) to find evolutionary equilibria and analyse their stabilities.

#### Model without the brake

We first study the properties of our model when the brake is absent (setting all variables containing the brake allele equal to zero). We determine the equilibrium states where only one allele is present (i.e. boundary equilibria). At the wild-type-only equilibrium, we have 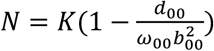, *p_D_* = 0, *δ*_0*D*_ = 0(see Mathematica Appendix for details). At the drive-only equilibrium, the size of the population depends on the type of drive. Since we only consider eradication drives here (i.e. drives such that a drive-only population is not viable), we have *N* = 0, *p_D_* = 1, *δ*_0*D*_ = 0 at the drive-only equilibrium (for completeness though, we included in the Mathematica appendix a separate stability analysis of the drive-only equilibrium for replacement drives). Generic formulas for interior equilibria (i.e. for which 0 < *p_D_* < 1) could not be found analytically.

#### Model with the brake

For simplicity, in the full model with the three alleles, we only study the stability of the wild-type-only equilibrium (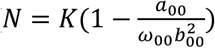, *p_D_* = 0, *p_B_* = 0, *δ*_0*D*_ = 0, *δ*_0*B*_ = 0, *δ*_*DB*_ = 0).

### Numerical solutions and stochastic simulations

#### Deterministic solutions of the model

To test the robustness of the equilibrium states predicted by our analytical model, we solve the model numerically for specific sets of parameters, using the original formulation in equation (1). We use parameter values for a threshold-independent eradication drive (i.e. as explained in the result section below, conditions where, according to the stability analysis of our model, the wild-type population cannot be recovered after the introduction of the brake). Time is discretized; we consider small fixed time steps *dt* = 0.005. When the system undergoes oscillations, genotype densities can go down to extremely small values, possibly below computer precision. We therefore set a critical value *thr* = 0.01, below which the density of a genotype is considered to be zero.

#### Stochastic simulations

To explore the effect of stochasticity on our model, we implement a stochastic version of it using a Gillespie algorithm (Gillespie 1977), directly translating the system of Ordinary Differential Equations (system (1) and the Appendix) into a stochastic simulation. In short, the algorithm goes as follows. Within a time step we (i) compute the rates (or “propensities”) of all possible events (birth and death probabilities of each of the five genotypes); (ii) randomly pick one event (the higher the event’s rate, the more likely its occurence); (iii) update the population according to the event that has taken place; (iv) draw the time interval that lasted the step (according to an exponential distribution parameterized by the sum of all propensities). For each set of parameter values, we run 10000 simulations, each of them until a maximum time value (*t_max_* = 25000) or until the population goes extinct. For each simulation, we list the different types of outcome (i.e., WT recovery after introduction of the brake, coexistence between the wild type and either the brake or both the initial gene drive and the brake, extinction before or after the introduction of the brake, drive loss before brake introduction).

### Data availability

Supplemental Material Files S1-S2 is available at Figshare: https://doi.org/10.6084/m9.figshare.11982285.v1

File S1 contains a supplemental script for the analytical model (Mathematica notebook). File S2 contains scripts for numerical explorations and stochastic simulations.

## Results

To assess the efficiency of various types of brakes to control gene drives, we use a combination of (i) analytical techniques (stability analysis of the deterministic model), (ii) numerical solutions of the deterministic model, and (iii) stochastic simulations. The stability analysis (i) is done with generic parameters. For the numerical steps of our exploration of the model ((ii) and (iii)), we use specific parameters corresponding to threshold-independent eradication drives, i.e. drives that spread from very low frequencies, and whose fixation leads to the extinction of the population.

### There are four categories of homing drives

To better understand the dynamics of the full model with three alleles (wild-type, drive, brake), we first study the model in the absence of brake. This analysis is done using generic parameters, separately for each timing of gene conversion (zygote vs. germline conversion).

In this two-allele version of the model, there are two boundary equilibria: drive loss (the wild-type allele is fixed) and drive fixation. These two equilibria can be locally stable or unstable, so that there are up to four possible combinations of stabilities and therefore four possible outcomes: (i) drive loss, (ii) coexistence of the drive and wild-type alleles, (iii) drive fixation, (iv) bistability (Deredec *et al*. 2008; Alphey and Bonsall 2014; Unckless *et al*. 2015; Noble *et al*. 2017; Vella *et al*. 2017; Girardin *et al*. 2019). Drives in (ii) and (iii) will invade the wild-type population from an arbitrary low frequency and are “threshold-independent” (Marshall and Akbari 2018). Drives in (iv) can either spread and fix when the drive allele is introduced at a high enough frequency or will be lost when their introduction frequency is below a given threshold (i.e. there is a bistability). This type of drive is “threshold-dependent” (Akbari *et al*. 2013; Marshall and Akbari 2018). The parameter ranges corresponding to each outcome are illustrated in Supplemental Material, Figures S2–S3, for replacement and eradication drives; they are consistent with the findings of previous studies (Deredec *et al*. 2008; Unckless *et al*. 2015; Vella *et al*. 2017; Girardin *et al*. 2019). The eradication drives used so far in laboratory studies (Kyrou *et al*. 2018) (large fitness cost, high conversion efficiency, recessivity and conversion in the germline) correspond to threshold-independent drives.

### Stability analyses indicate that a brake can recover the wild-type population only if the drive is threshold-dependent

When the brake allele has lower fitness than the wild-type allele, the wild-type, drive and brake alleles, are involved in non-transitive interactions (rock-paper-scissors type; Vella *et al*. 2017): the wild-type is converted into a drive by the drive, the drive is converted into a brake by the brake, and the brake is costly compared to the wild-type. A high frequency of the wild-type, drive or brake in the population favors the drive, brake or wild-type respectively. Such negative frequency-dependent selection can result in the coexistence of the three alleles.

In the analytical model with the three alleles, we find that the conditions for the local stability of the wild-type-only equilibrium are the same as in the model without brake (details of the calculations are presented in the supplementary Mathematica file). In other words, our stability analysis indicates that the introduction of a brake can successfully restore a wild-type population only under two conditions. First, quite trivially, the wild-type population can be recovered when the population is targeted by a drive that would be lost in the absence of brake (drive loss equilibrium above; we ignore this case thereafter). Second, the wild-type population can be recovered when it is targeted by a threshold-dependent drive (i.e. with parameters corresponding to a bistability in the model without brake, see above). In this case, introducing the brake allele can decrease the frequency of the drive allele below its invasion threshold; the drive is then lost. Once the drive is lost, if it is, the brake loses the competition against the wild-type allele because of its fitness cost, and the wild-type population is finally recovered.

### Numerical explorations of the deterministic model and stochastic simulations show that brakes can stop threshold-independent drives under certain conditions

#### Numerical solutions of the deterministic model

The introduction of a brake in a population targeted by a threshold-independent drive may lead to oscillations of large amplitude. During these oscillations, the densities of some genotypes may reach extremely low values. Analytically, no allele should get lost in these oscillations because we assumed infinite population sizes in the analysis. Biologically, this is not realistic: however big a population, an extremely low density may correspond to less than one individual, and thus to the loss of an allele from the population. Computationally as well, these oscillations are challenging, because they may lead to values below the minimum number that a computer can represent, and therefore to the failure of numerical solutions. To solve both issues, we set a critical density below which a genotype is considered absent from the population and we numerically integrate our model to further explore the effect of the introduction of a brake in a population targeted by a threshold-independent eradication drive. Cutting large amplitude cycles means that alleles can be lost. The dynamics of the frequencies of the three alleles and of population size (scaled by the equilibrium density of the wild-type population) are shown in Figure 2. These dynamics depend on the trait that is affected by the drive and the brake (fecundity, adult mortality, or zygote survival; lines in Figure 2), the level of dominance (columns in Figure 2), and whether the brake restores fitness or not (Supplemental Material, Figures S4 vs. Figure 2).

**Figure 2:**
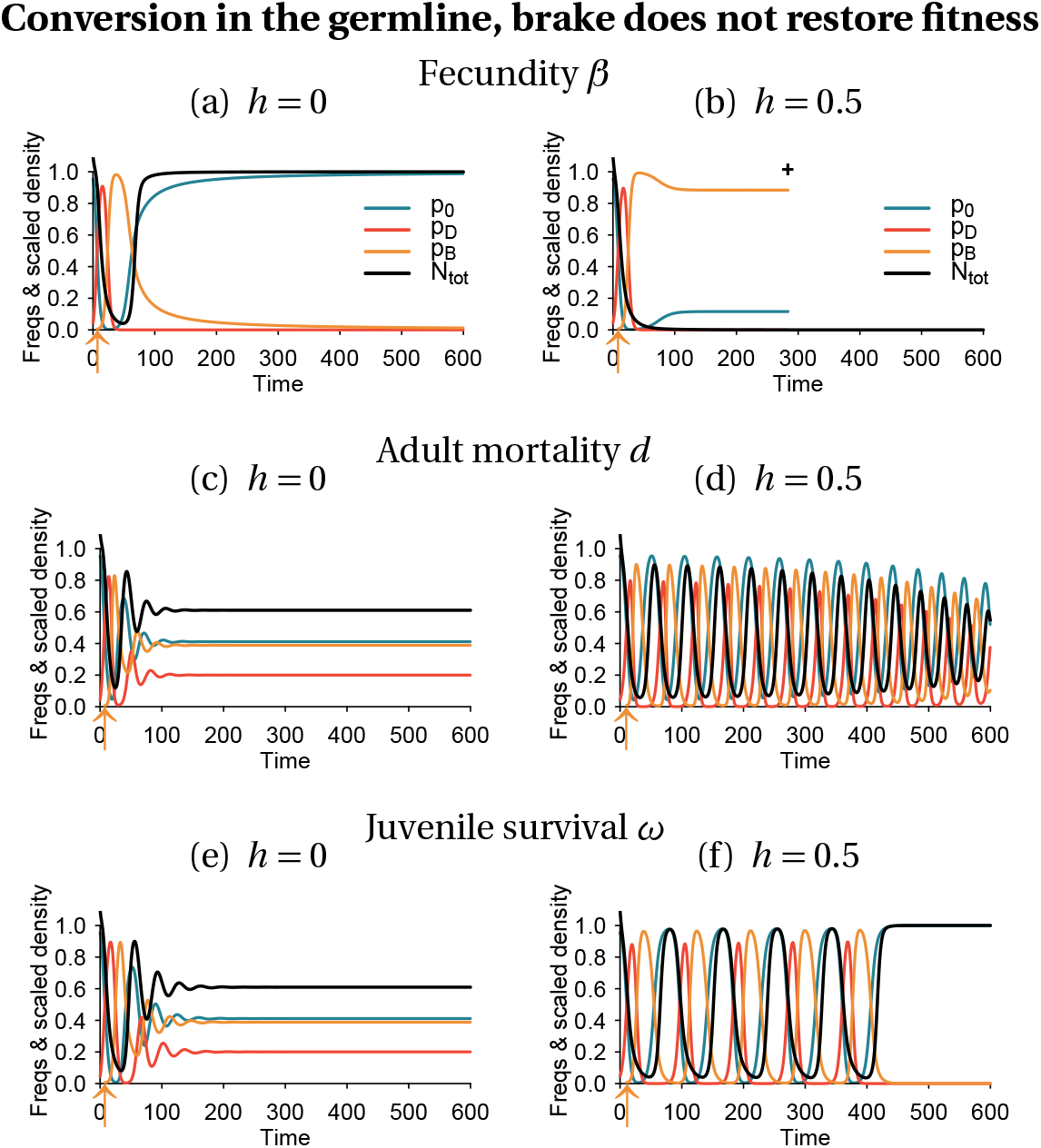
Deterministic dynamics of the frequencies of each allele in the population, and scaled total population size (black curve). Conversion takes place in the germline, and the brake does not restore fitness. Population size is scaled relative to the equilibrium size of a 100% wild-type population 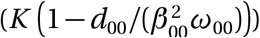. The arrow indicates the timing of drive introduction, here chosen to be when the drive allele is at 50% (*f_I_* = 0.5). A cross indicates population extinction. Simulation parameters are listed in Tables S1–S3.

The addition of a critical minimum density leads to outcomes that were not predicted by our stability analysis. Contrary to the predictions of the stability analysis for threshold-independent drives, in Figures 2(a) and 2(f), the drive is lost, allowing for population recovery. This is because the density of drive-carrying individuals reaches so small values at some point that the drive allele is considered extinct. Then, the brake allele being costly compared to the wild-type allele, it decreases in frequency and is lost as well. In Figure 2(b), the population goes extinct. This is because the overall population density goes down to very small values.

As expected, with our parameters, the wild-type population is more rarely recovered with a brake that does not restore fitness than with a brake that does (compare Figures 2 to S4, and S5 to S6).

We hypothesized that allele loss would happen when the amplitude of oscillations increases (i.e. when the interior equilibrium, where the three alleles coexist, is unstable). However, even when the amplitude of oscillations decreases (i.e. when the interior equilibrium is locally stable), the initial oscillations can be substantial, hindering our ability to predict the outcome. In addition, the outcome itself depends on non-biological contingencies such as the time interval at which the solutions are calculated and the critical density below which a genotype is considered extinct. As a consequence, a brake is not guaranteed to prevent the eradication of a population targeted by a threshold-independent drive.

#### Stochastic simulations

We complemented our exploration with stochastic simulations. Notably, having integer numbers of individuals of each genotype avoids the arbitrary choice of a critical density below which a genotype is considered extinct. Importantly, the parameters that we chose in our simulations correspond to a large wild-type population size (an expected density of N* = 10000); the diversity of observed outcomes is due to the large amplitude of oscillations in genotype densities triggered by the introduction of the brake.

Among the different parameters investigated, whether or not the brake restored fitness has the highest impact on the recovery of the wild type population (Figure 3 vs. 4 and 5 vs. 6). Our stochastic simulations show that in many instances, the brake does not prevent population extinction when it does not restore fitness (Figures 3 and 5). In contrast, the drive allele is always lost when the brake restores fitness (Figures 4 and 6), resulting either in the full recovery of the wild-type population, or in a coexistence between the wild type and the brake at the time at which the simulation ended (tmax = 2500). Noteworthily, as the fitness of the brake approaches that of the wild-type allele, the time necessary to recover 100% wild-type individuals increases.

**Figure 3:**
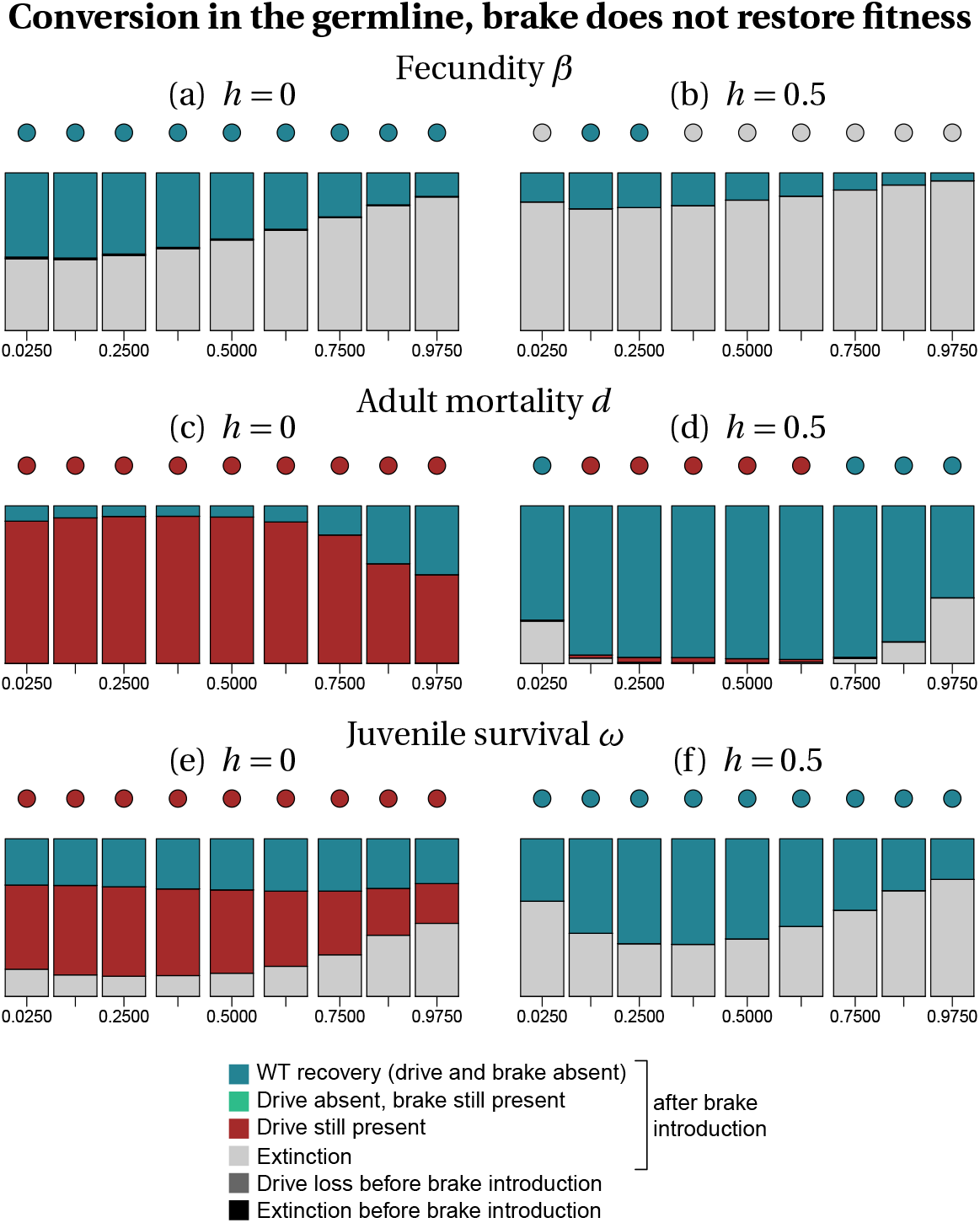
Frequency of each type of outcome in the simulations (color-coded), depending on the frequency of drive *f_I_* at the time at which the brake is introduced (horizontal axis), on the dominance coefficient *h* (columns) and on the trait that is affected by the drive and the brake (rows). The dots show, with the same color code, the output of the deterministic model. Simulation parameters are listed in Tables S1–S3.

**Figure 4:**
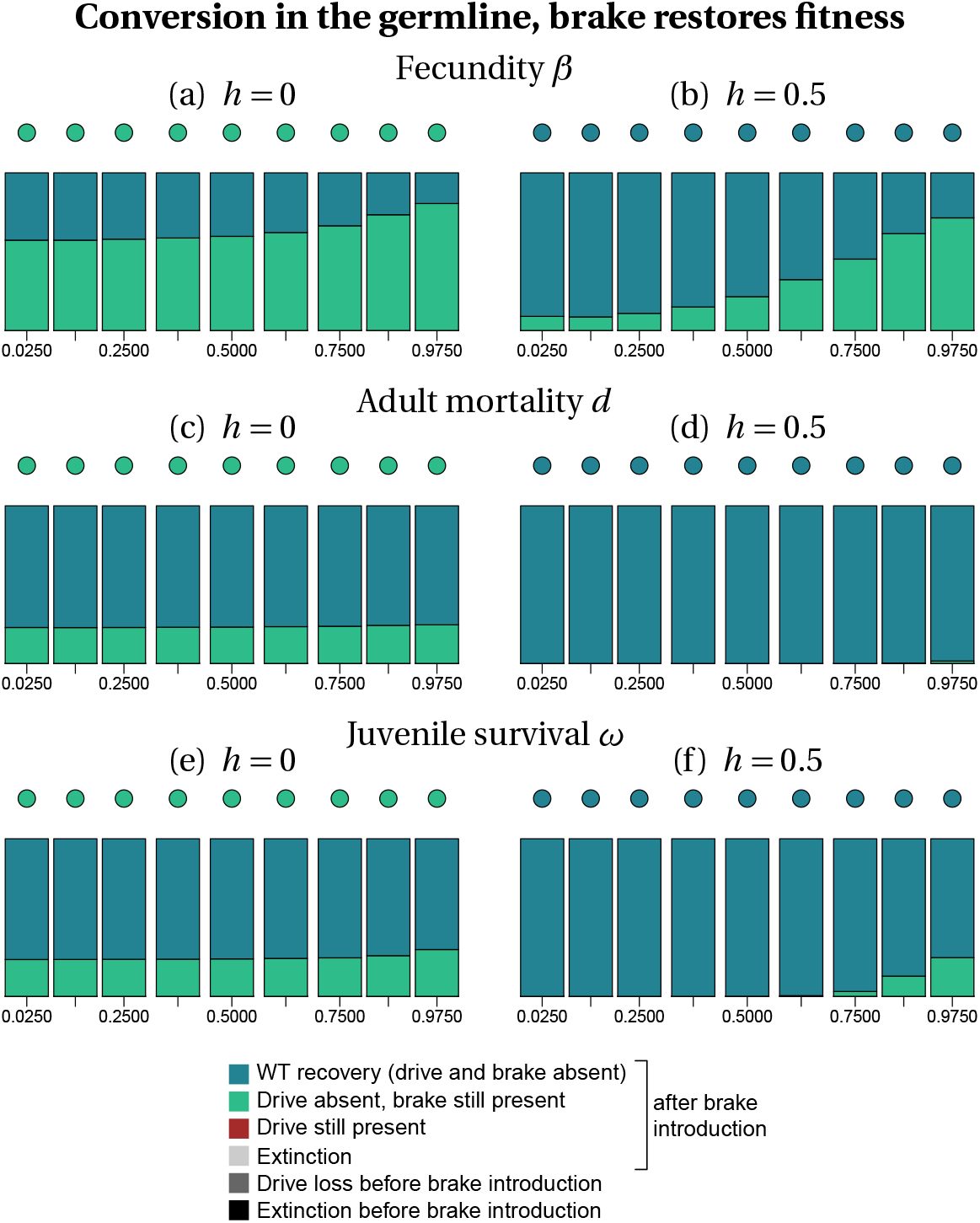
Frequency of each type of outcome in the simulations (color-coded), depending on the frequency of drive *f_I_* at the time at which the brake is introduced (horizontal axis), on the dominance coefficient *h* (columns) and on the trait that is affected by the drive and the brake (rows). The dots show, with the same color code, the output of the deterministic model. Simulation parameters are listed in Tables S1–S3.

**Figure 5:**
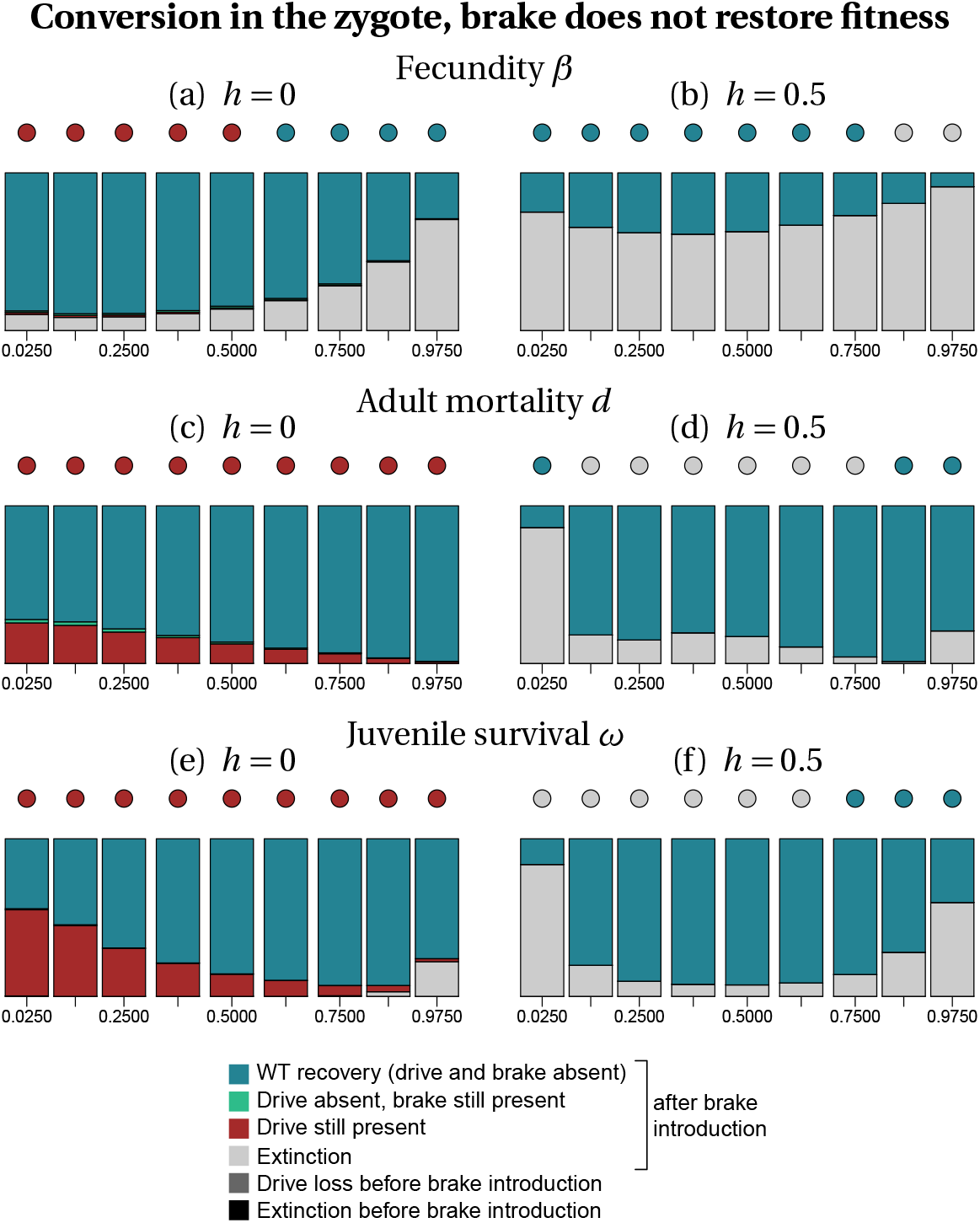
Frequency of each type of outcome in the simulations (color-coded), depending on the frequency of drive *f_I_* at the time at which the brake is introduced (horizontal axis), on the dominance coefficient *h* (columns) and on the trait that is affected by the drive and the brake (rows). The dots show, with the same color code, the output of the deterministic model. Simulation parameters are listed in Tables S1–S3.

**Figure 6:**
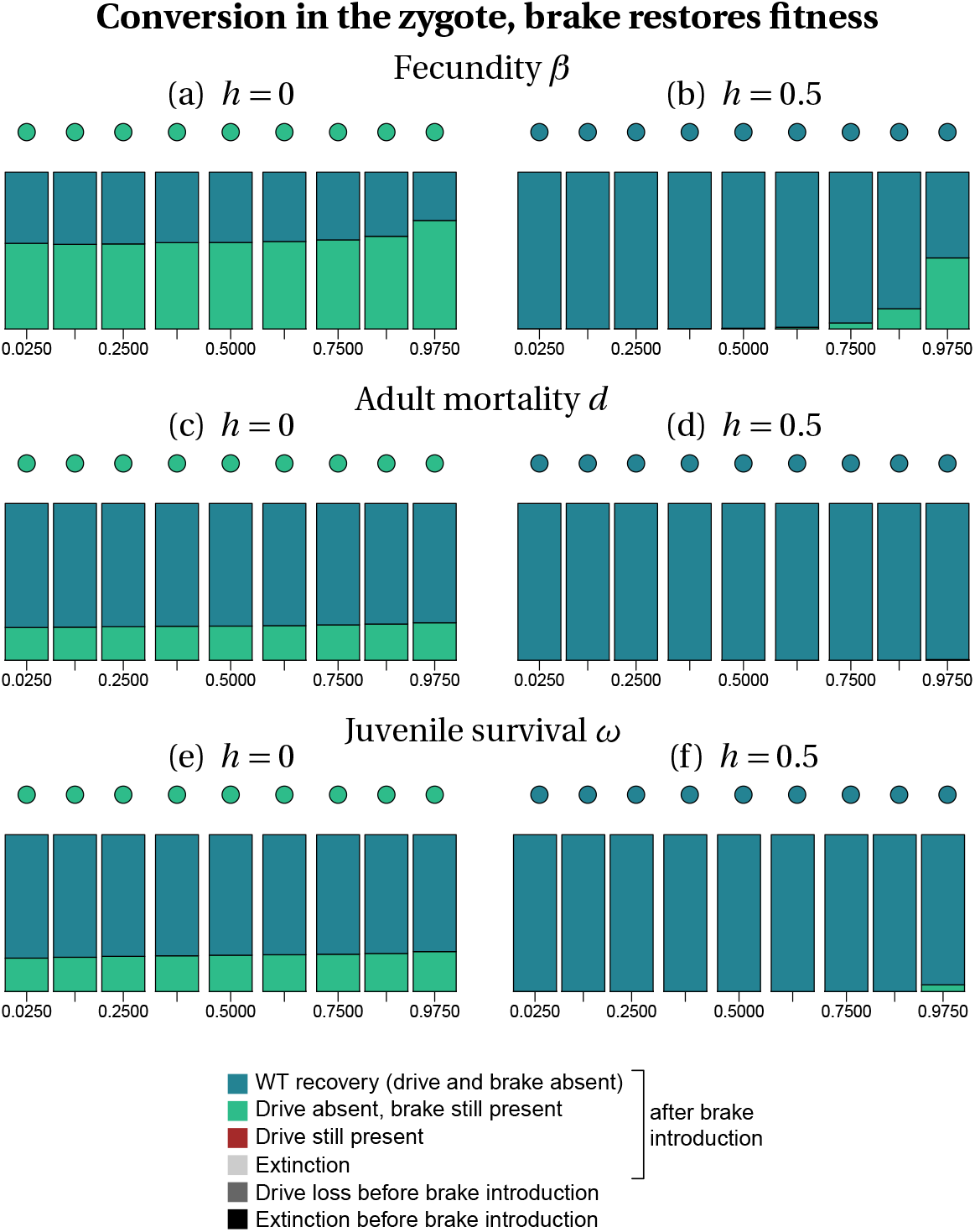
Frequency of each type of outcome in the simulations (color-coded), depending on the frequency of drive *f_I_* at the time at which the brake is introduced (horizontal axis), on the dominance coefficient *h* (columns) and on the trait that is affected by the drive and the brake (rows). The dots show, with the same color code, the output of the deterministic model. Simulation parameters are listed in Tables S1–S3.

When the brake does not restore fitness, the recovery of the wild-type population is more frequent when gene conversion occurs in the zygote than when it occurs in the germline, especially for recessive drives and brakes (*h* = 0, Figure 3 vs. 5). When the brake restores fitness, the timing of conversion has little effect on the final outcome (compare Figure 4 with Figure 6). The likelihood of recovering a 100% wild-type population often decreases with drive frequency at brake introduction, i.e. with later brake introductions. Early brake introductions (i.e. introductions when the drive frequency is still low) may nevertheless fail, for instance due to stochastic loss of the brake. The effects of other parameters such as the type of trait targeted or the level of dominance are more difficult to predict. The most frequent outcome in stochastic simulations was often different from the outcome predicted by deterministic models. For example, population extinction is the most frequent outcome of some of the stochastic simulations, while the corresponding deterministic model predicts the recovery of the wild-type population (e.g. Figures 3(a), 5(b)). We conclude, in agreement with the results of Vella *et al*. (2017) using infinite population size, that a brake is not guaranteed to prevent the eradication of a population targeted by a threshold-independent eradication drive.

## Discussion

We developed a model to investigate the consequences of introducing a brake allele in a population targeted by a CRISPR-based homing gene drive. In contrast to previous models that assumed 100% cleavage efficiency in the germline and only considered threshold-independent gene drives (Vella et al. 2017; Girardin et al. 2019), our model also considers imperfect cleavage and threshold-dependent gene drives. Our framework also extends previously published models, which focused on allele frequencies (ignoring fluctuations in population density, Vella *et al*. 2017; Girardin *et al*. 2019). By accounting for the effects of both the initial gene drive and the brake on population size, our model represents a first step towards the explicit integration of changes in population size into the prediction of the dynamics of wild-type, gene drive and brake alleles. While we concentrate here our numerical explorations on eradication drives and threshold-independent drives, our model can also be used to study the dynamics of replacement drives and their brakes, by adapting parameter values. Our model can form a basis for future studies investigating the effect of CRISPR-based brakes against other types of gene drives (e.g. split gene drives; Li *et al*. 2020), to check whether these alternatives might be easier to control.

Our model does not account for the potential evolution of resistance against gene drives. Such resistance can be due to cleavage repair by non-homologous end joining or to natural variation at the target locus, and can occur frequently after the release of gene drive individuals (Drury *et al*. 2017; Unckless *et al*. 2017; Bull *et al*. 2019). However, several strategies are under way to prevent the evolution of gene drive resistance, such as the use of multiple gRNAs (Champer *et al*. 2018; Oberhofer *et al*. 2018; Edgington *et al*. 2020) or the targeting of a functionally constrained locus whose mutations are highly deleterious and cannot increase in frequency (e.g. Kyrou *et al*. 2018). Given these efforts to limit the evolution of resistance against gene drives, we chose not to include this feature in our model. In addition, Vella et al. (2017) investigated the evolution of resistance at the target locus in addition to the introduction of a countermeasure and found that the qualitative behavior of the brake remains unchanged (polymorphic equilibrium of all alleles).

Furthermore, we did not model the evolution of resistance against brakes either. Developing new brake constructs to counter resistance would be both costly and time consuming, so that developing brakes that are the least sensible to the evolution of resistance is important. So far only CATCHA brakes have been developed in the laboratory (Wu et al. 2016). If resistant alleles were to form, for the types of brakes we investigated, the consequences would differ between ERACR and CATCHA brakes. For ERACR brakes, mutations arising in flanking sequences targeted by the brake could prevent cleavage and conversion of the drive into a brake. If these mutations do not alter the rate of conversion of the wild-type allele into a drive allele, a drive resistant to the ERACR brake could continue spreading. Thus, ERACR brake could fail to prevent a population from extinction. For CATCHA brakes, mutations in the target *cas9* sequence would result in non-functional Cas9 enzymes. These brake-resistant alleles would have the same fitness cost as the drive allele, but without the gene-conversion advantage of the drive. Should they appear, they would be expected to remain at a low frequency in the population. Overall, we thus expect CATCHA brakes to overcome the evolution of resistance against brake while ERACR brakes would not, so we recommend using the former.

Overall, our model shows that the success of recovering the wild-type population using a brake depends both on the type of brake introduced and the type of gene drive targeted. More specifically, our conclusions depend on the method chosen to explore the model. Our stability analysis indicates that the wild-type population can only be recovered after the introduction of a brake if the drive is threshold-dependent. Nevertheless, our numerical integration of the model -- including a critical population density to avoid unrealistically low genotype densities -- and stochastic simulations show that the wild-type population can also be recovered in certain cases when a threshold-independent drive is used. In these cases, brakes that restore fitness can better control a gene drive than universal brakes that do not. However, we could not draw general conclusions on the effect of other parameters (e.g. fitness trait affected by the drive, dominance level, timing of conversion, and frequency of the drive for introducing the brake) on the final outcome.

Our model shows that, even when the brake is introduced when the eradication drive is still at a low frequency, the frequency of the eradication drive continues to increase and results in a strong population bottleneck (e.g. Figure 2a). Such a strong bottleneck could result in a long term alteration of the recovered wild-type population (e.g. due to inbreeding depression). This point is important to keep in mind even though it is not explicitly incorporated in our model.

Our study has practical implications. First, we advise against using universal brakes as the sole countermeasure because they are not guaranteed to succeed and stop a drive. In contrast, we recommend using specific brakes which include a recoded version of the gene disrupted by the initial gene drive. Since they restore fitness, they are more likely to be effective: they spread at a faster rate and increase the chances of recovering a population of wild-type individuals. To reduce potential environmental risks, we recommend that the development of homing gene drives goes in pair with the co-development of such specific brakes. Although they are not guaranteed to successfully restore a 100% wild-type population, specific brakes currently represent the best countermeasure against the spread of homing drives following an escape from a laboratory. We also recommend laboratory studies to assess the efficacy of brakes using experimental evolution under controlled conditions. Second, because they are easier to control with brake, we believe that threshold-dependent homing gene drives are a safer alternative to threshold-independent homing drives, that are currently being developed in laboratories. These threshold-independent homing drives are characterized by large and recessive large fitness costs, high conversion efficiency and germline conversion (e.g. Kyrou *et al*. 2018). Several studies (Tanaka *et al*. 2017; Min *et al*. 2018) have recommended the use of spatially and/or temporally limited threshold-dependent homing drives, because they are less likely to spread into non-target populations. However, we emphasize that it might be difficult in practice to implement a threshold-dependent drive whose threshold remains as expected for several reasons. First, theoretical models show that the range of parameter values for threshold-dependent gene drives is larger when conversion occurs in the zygote than when it occurs in the germline (compare Figures 1 and 4 in Deredec *et al*. 2008; Figure S2–S3). So ideally, it might be better to use drives with conversion in the zygote. Nevertheless, such drives are more difficult to create and so far all successful homing drives have been engineered with germline promoters (Table 2 in Courtier-Orgogozo *et al*. 2019b). A few conserved genes are expressed in the germline of all animals (*nanos, vasa, piwi*; Extavour and Akam 2003; Juliano *et al*. 2010) and their promoters constitute preferred tools for engineering gene drive constructs in various animal species, in contrast to zygotically expressed genes, which tend to be less conserved across taxa (Heyn *et al*. 2014). Second, “real life” ecological conditions are likely to alter the genetic parameters of any gene drive, in particular its fitness cost. Fitness costs are difficult to estimate in the field and can vary either across genomic backgrounds, spatially or temporally (Marshall and Hay 2012; Backus and Delborne 2019). Hence, depending on ecological conditions, the threshold value for the invasion of a threshold-dependent homing drive could change, or even decrease to 0. Thus, a homing drive that is threshold-dependent in the laboratory might turn into a threshold-independent drive in the wild.

### Conclusion

Our model is a step towards the development of more complex analytical models of gene drive that account for the feedback between population demography and evolution. Our results suggest that the recessive eradication drives with germline conversion currently developed in mosquitoes (e.g. Kyrou *et al*. 2018) are likely to be threshold-independent and could be particularly difficult to control using brakes. In addition, our results show that a brake that carries a version of the gene disrupted by the initial gene drive, and therefore restores fitness, can prevent the extinction of the target population under certain conditions. We recommend that the development of countermeasures should go in pair with the development of drives. Given the diversity of outcomes that we find and the difficulty to precisely estimate the relevant parameters determining each outcome, specific experimental studies will be necessary to confirm modelling outcomes that a given brake can indeed stop the spread of drives. A brake should not be considered reliable before population experiments are carried out.

## Acknowledgements

We thank Arnaud Martin for discussions. Funding for this project was provided by European Research Council (FP7/2007-2013 Grant Agreement no. 337579) to VCO, ANR-14-ACHN-0003 and ANR-19-CE45-0009-01 to FD and the CeMEB LabEx/University of Montpellier (ANR-10-LABX-04-01) and an INRAE-SPE starting grant to NOR.

## Author contributions

VCO brought the research topic, all authors developed the model, FD did the analysis, implemented numerical solutions, ran stochastic simulations. All authors analysed data and wrote the manuscript.

## Appendix

In the main text, the change over time in the density of individuals of genotype *g* is given by

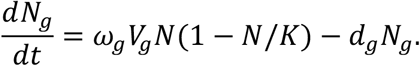

We provide below the expressions for *V_g_* for the two timings of gene conversion that we consider in the article.

### Germline conversion

When gene conversion takes place in the germline, individuals born heterozygous remain heterozygous as adults, their life-history parameters are those of heterozygotes, but then gene conversion takes place in the germline, and if successful, predominantly one type of gamete is produced by the individual. We have

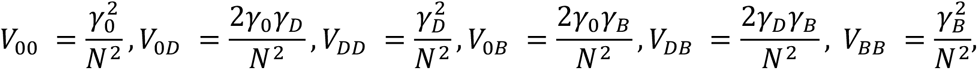

where

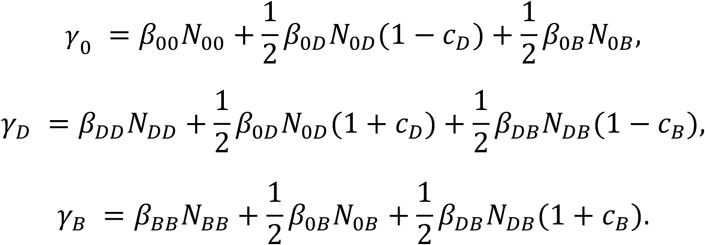

### Zygote conversion

When conversion takes place in zygotes, and when gene conversion is successful, an initially heterozygous zygote becomes homozygous, and develops into a homozygous adult. We have

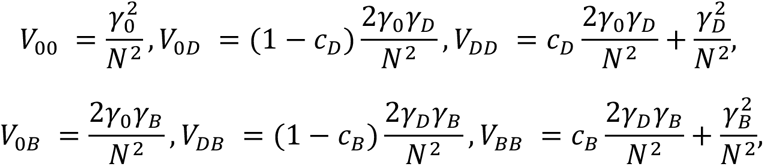

where

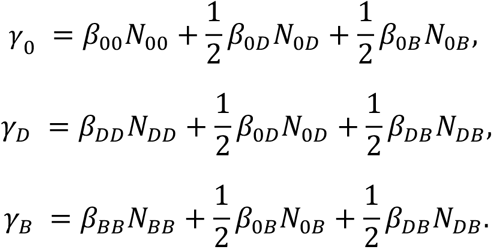

### Hypotheses regarding dominance

Here we justify why we can consider that the dominance parameter *h* is the same for all alleles. Let us first assume that the brake allele does not restore fitness. Under this scenario, the brake and gene drive alleles are genetically equivalent so that they have the same fitness (*ω_DD_* = *ω_BB_*) and the same dominance (*ω*_0*D*_ = *ω*_0*B*_). This is consistent with having the same dominance parameter:

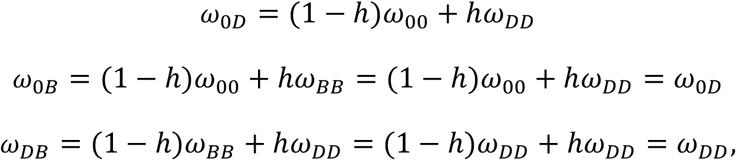

and likewise for *d* and *β* parameters.

Now let us assume that the brake allele does restore fitness. Under this scenario, the brake and wild-type alleles are genetically equivalent so that they have the same fitness (*ω*_00_ ~ *ω_BB_*) and the same dominance (*ω*_0*D*_ ~ *ω_DB_*). This is also consistent with having the same dominance parameter::

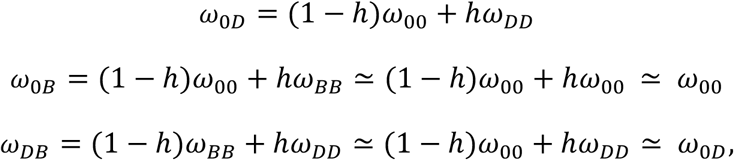

and likewise for *d* and *β* parameters. Therefore we can assume that dominance levels are equal across the three different types of heterozygotes both when the brake does and does not restore fitness.

---

**Table S1:** Fixed parameters

|  |  |  |
| --- | --- | --- |
| $K$ | Carrying capacity | 25000 |
| $c_D$ | Probability of gene conversion by a drive | 0.9 |
| $c_B$ | Probability of gene conversion by a brake | 0.8 |
| $N_{0D}^{(0)}$ | Initial number of introduced drive-WT individuals | 1000 |
| $N_{0B}^{(0)}$ | Initial number of introduced brake-WT individuals | 100 |
| $t_{\max}$ | Maximum time of the simulations | 2500 |

**Table S2:** Varying parameters

|  |  |  |
| --- | --- | --- |
| $f_I$ | Frequency of the drive allele in the population when the brake is introduced | {0.025, 0.1375, 0.25, 0.375, 0.5, 0.625, 0.75, 0.8625, 0.975} |
| $h_{D0} = h_{B0} = h_{DB} = h$ | Dominance parameter | {0, 0.5} |

**Table S3:** Parameters for the different scenarios, depending on whether the brake restores fitness (modulo a small cost) or not, and on which life-history parameter is affected (adult survival *d*, zygote survival *ω*, adult fecundity *β*).

| Scenario # |  | (1) | (2) | (3) | (4) | (5) | (6) |
| --- | --- | --- | --- | --- | --- | --- | --- |
| Brake... |  | does not restore fitness |  |  | restores fitness |  |  |
| Effects on | | $d$ | $\omega$ | $\beta$ | $d$ | $\omega$ | $\beta$ |
| Adult death rate | $d_{00}$ | 0.6 | 0.6 | 0.6 | 0.6 | 0.6 | 0.6 |
| | $d_{DD}$ | 1.1 | 0.6 | 0.6 | 1.1 | 0.6 | 0.6 |
| | $d_{BB}$ | 1.1 | 0.6 | 0.6 | 0.64 | 0.6 | 0.6 |
| Juvenile survival | $\omega_{00}$ | 1.0 | 1.0 | 1.0 | 1.0 | 1.0 | 1.0 |
| | $\omega_{DD}$ | 1.0 | 0.545 | 1.0 | 1.0 | 0.545 | 1.0 |
| | $\omega_{BB}$ | 1.0 | 0.545 | 1.0 | 1.0 | 0.938 | 1.0 |
| Adult fecundity | $\beta_{00}$ | 1.0 | 1.0 | 1.0 | 1.0 | 1.0 | 1.0 |
| | $\beta_{DD}$ | 1.0 | 1.0 | 0.738 | 1.0 | 1.0 | 0.738 |
| | $\beta_{BB}$ | 1.0 | 1.0 | 0.738 | 1.0 | 1.0 | 0.968 |

**Figure S1:**
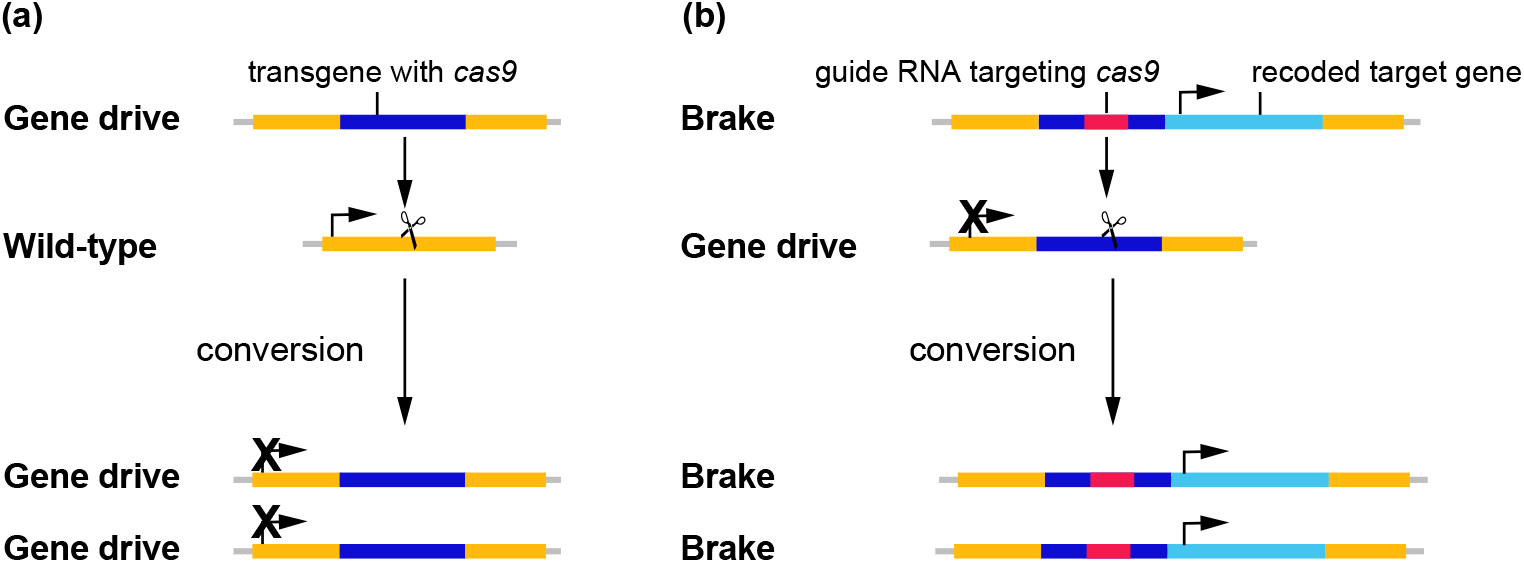
Gene conversions: (a) Conversion of the wild-type allele into a gene drive allele and (b) conversion of the gene drive allele into a brake allele that restores fitness. The brake construct includes a functional version (light blue) of the target gene (light orange) disrupted by the gene drive.

**Figure S2:**
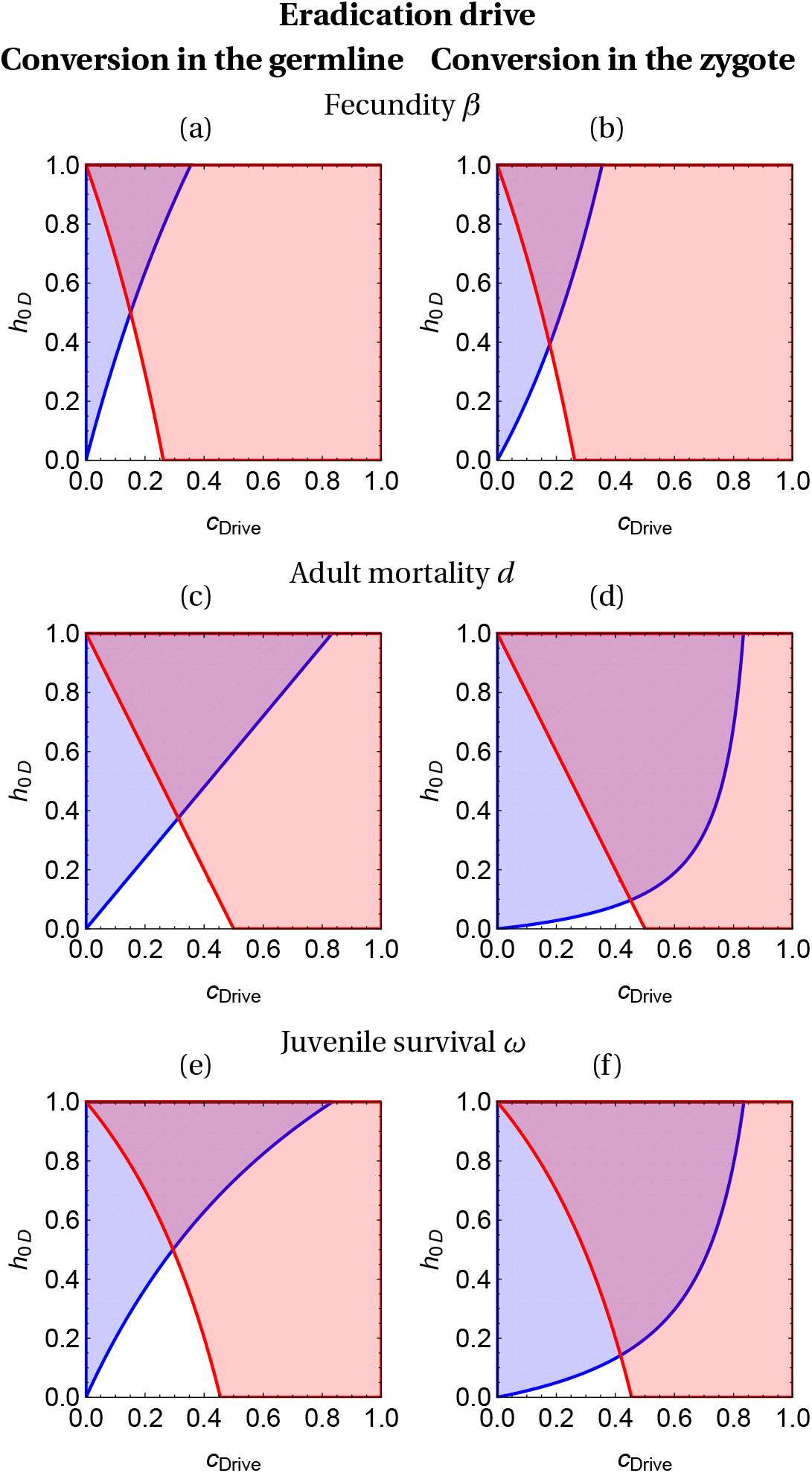
Local stabilities of the drive-only and the wild-type only equilibria in the absence of brake, for an eradication drive. The wild-type only equilibrium is locally stable in the blue-shaded region left of the blue curve; the drive-only equilibrium is locally stable in the red-shaded region right of the red curve. Neither equilibrium is locally stable in the white area, in which the two alleles coexist. Both equilibria are locally stable in the purple area; the final outcome depends on the initial conditions (bistability). Drives whose parameters put them in the purple area are threshold-dependent. Parameters are listed in Tables S1–S3.

**Figure S3:**
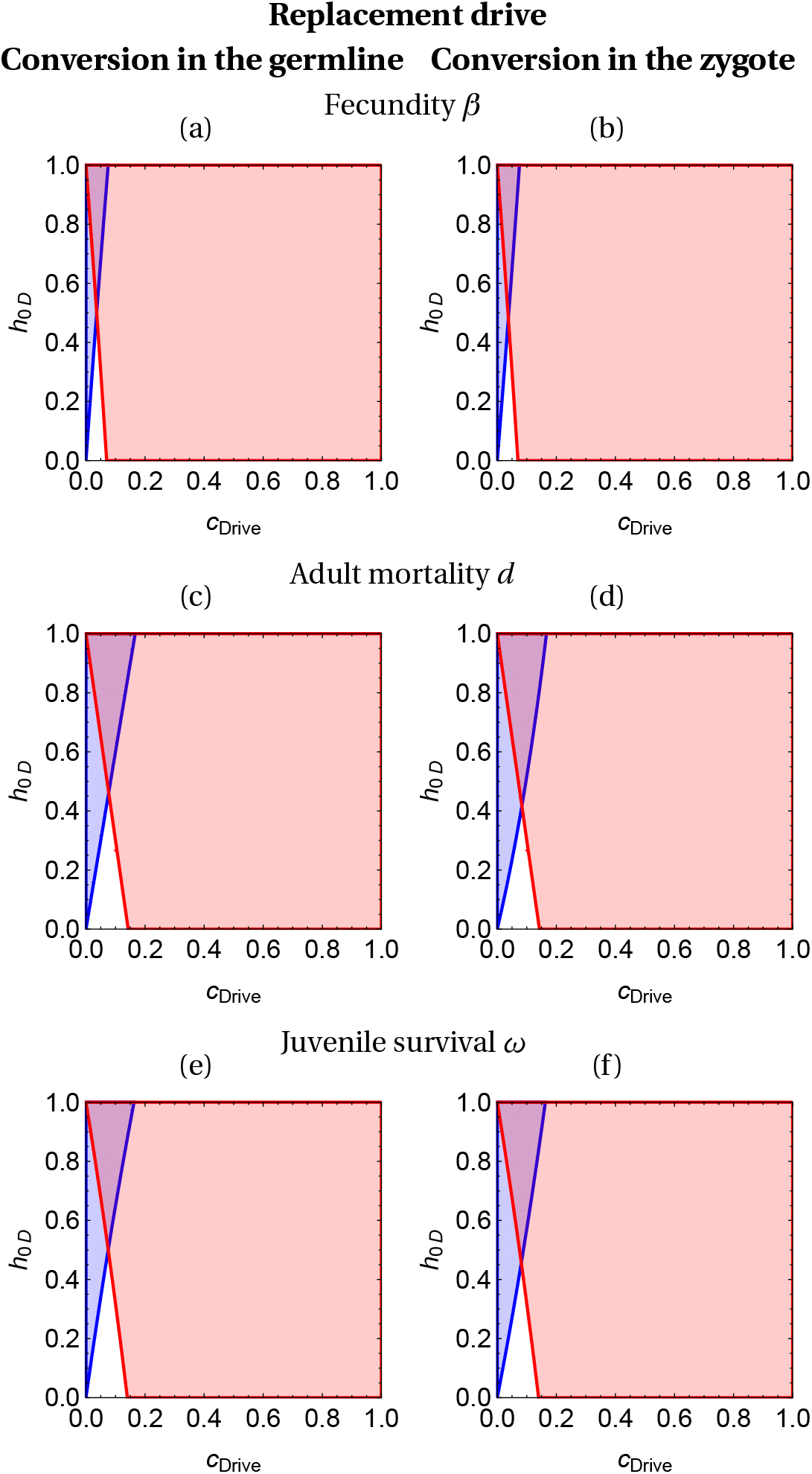
Local stabilities of the drive-only and the wild-type only equilibria in the absence of brake, for a replacement drive. The legend is the same as figure S2.

**Figure S4:**
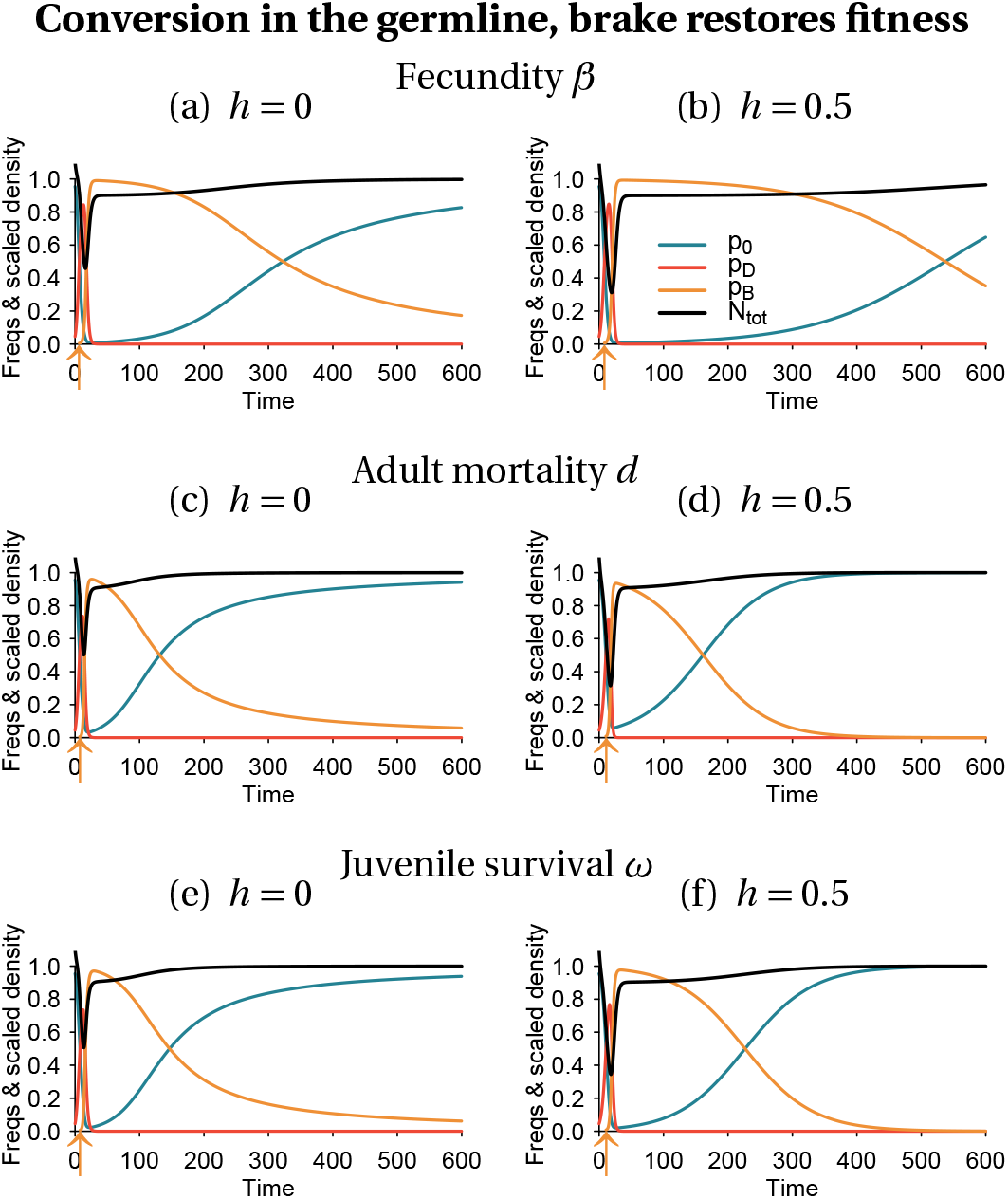
Same legend as figure 2.

**Figure S5:**
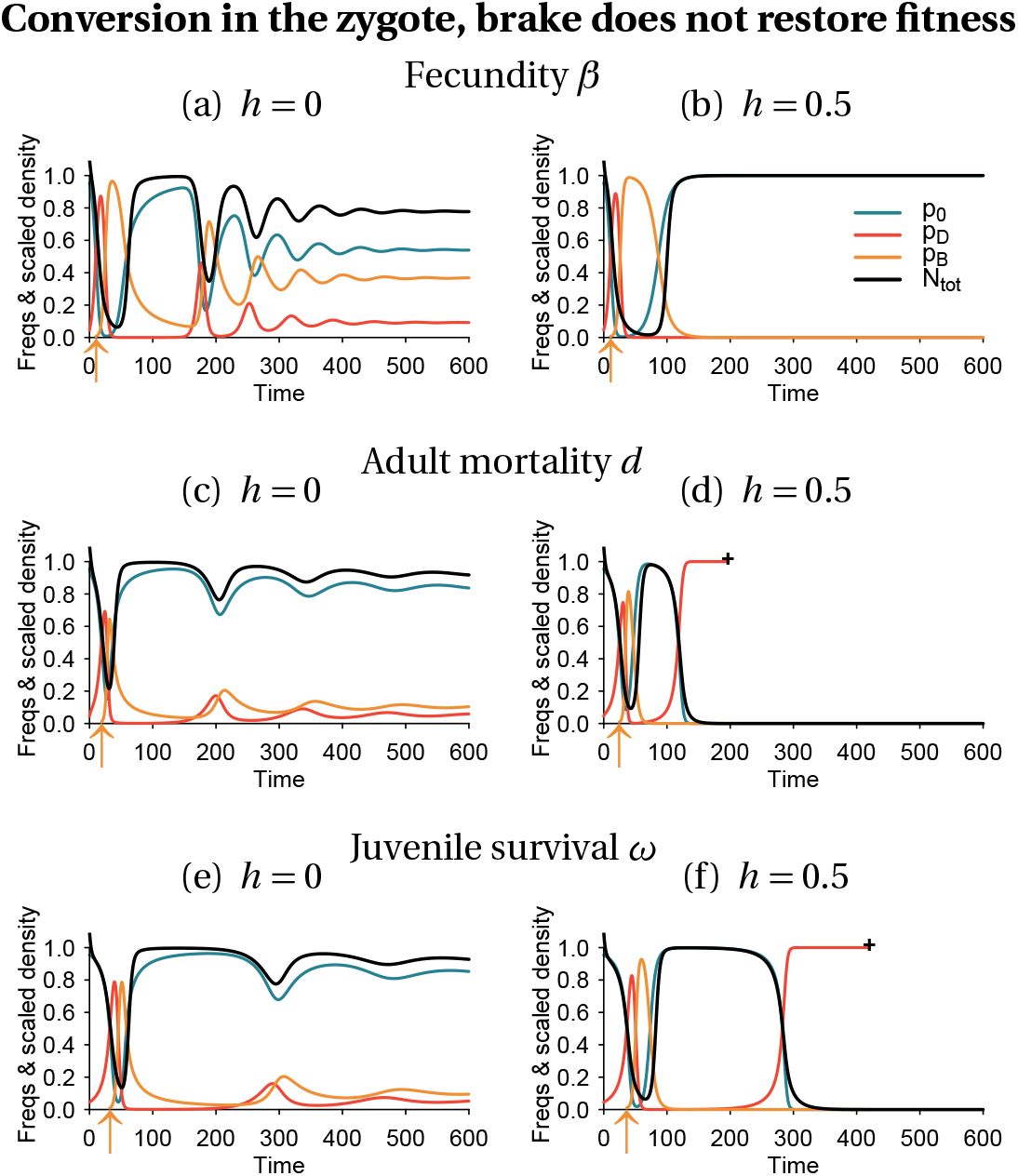
Same as figure 2

**Figure S6:**
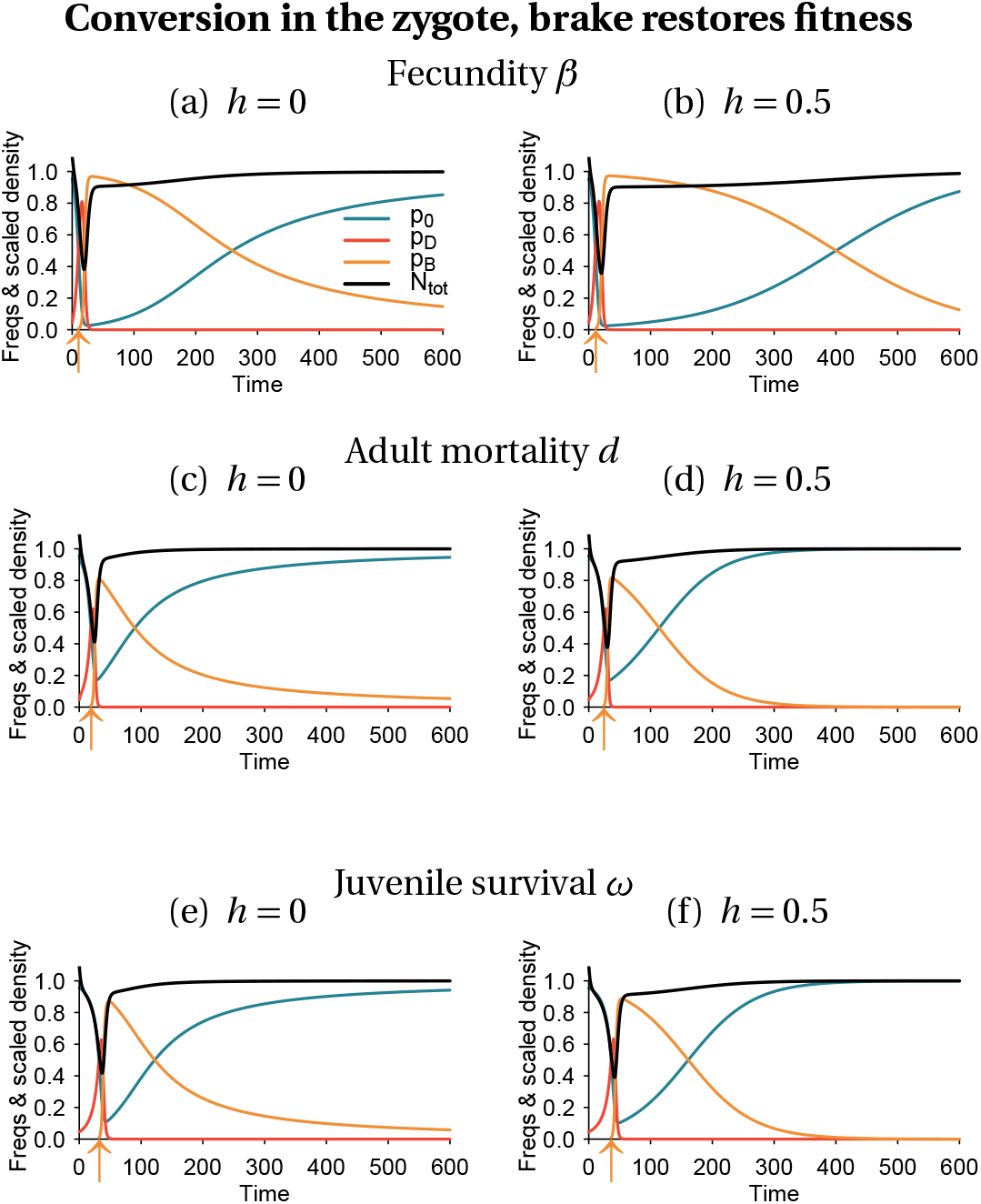
Same as figure 2

**Figure S7:**
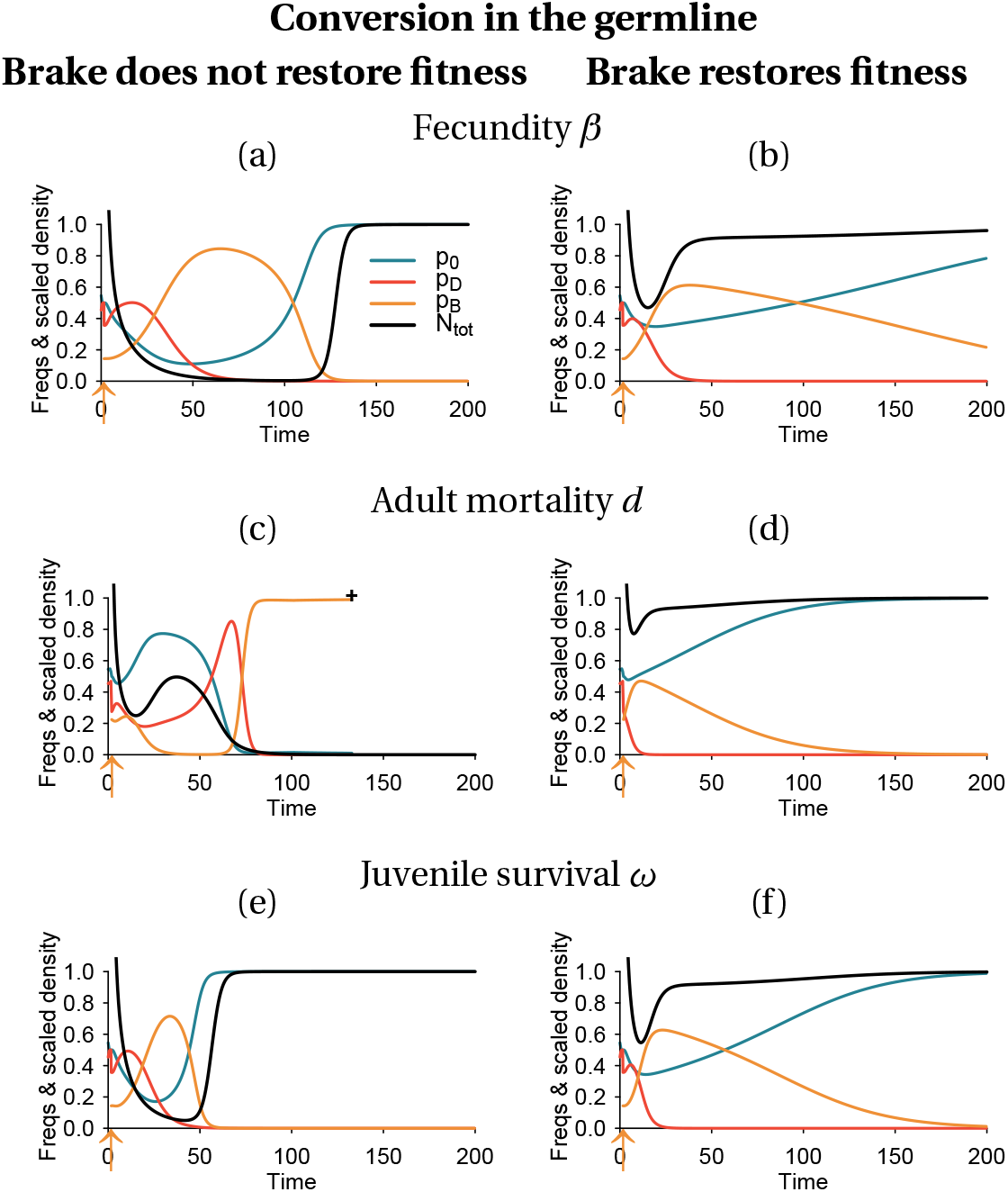
Deterministic dynamics when the drive is threshold-dependent; conversion takes place in the germline. Parameters are the same as in the other figures, except for the dominance parameter (*h* = 1) and for conversion efficiencies (*c*_D_ = 0.3, *c*_B_ = 0.25 in panels (a)–(b);*c*_D_ = 0.6, *c*_B_ = 0.55 in panels (c)–(d); *c*_D_ = 0.5, *c*_B_ = 0.45 in panels (e)–(f)). Introduction densities are *N*_0D_ = 10^5^ and *N*_0B_ = 10^4^.

**Figure S8:**
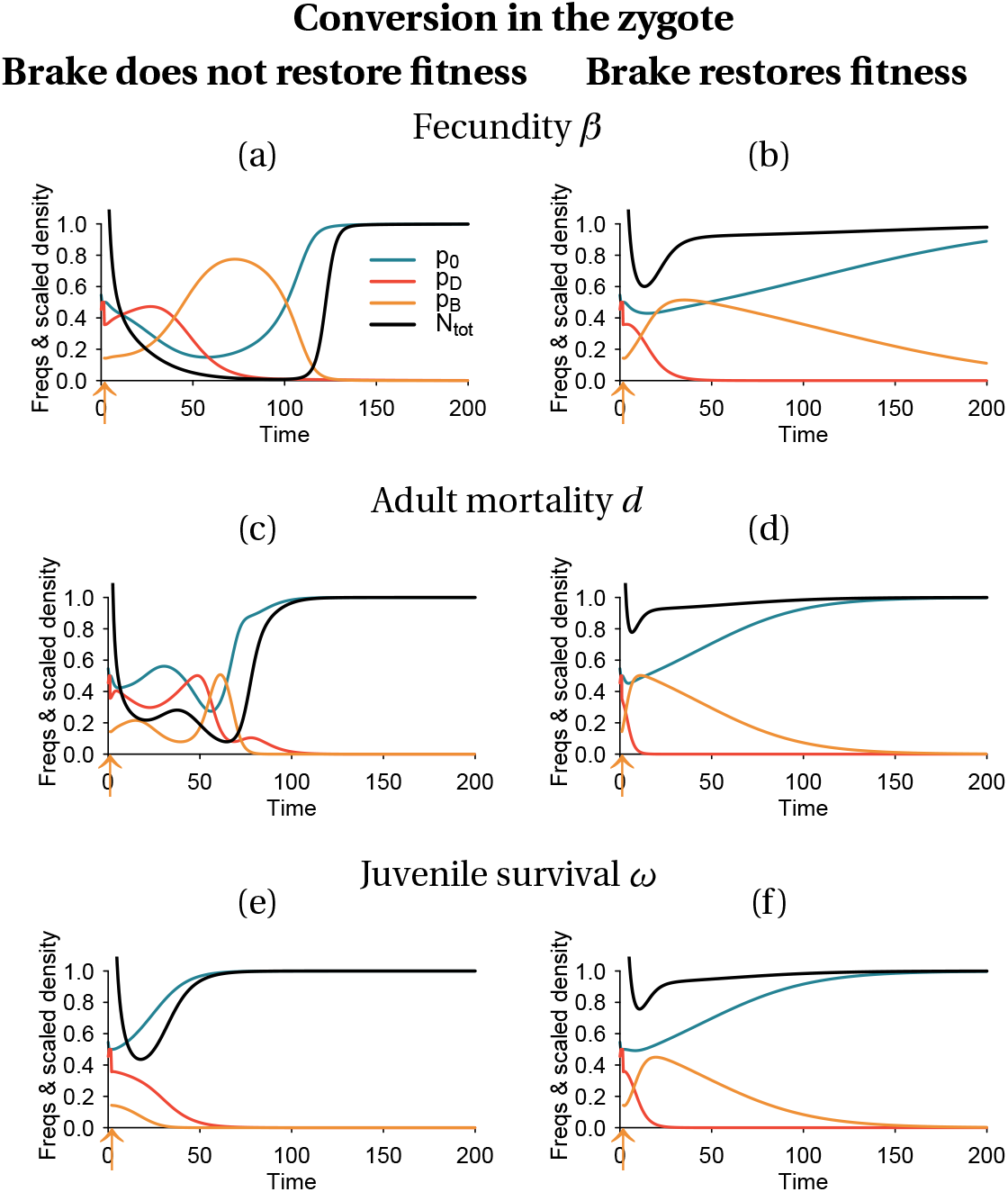
Deterministic dynamics when the drive is threshold-dependent; conversion takes place in the zygote. See figure S7 for parameter values.

